# Prediction of Individual Breast Cancer Evolution to Surgical Size

**DOI:** 10.1101/2020.06.13.150136

**Authors:** Cristian Axenie, Daria Kurz

## Abstract

Modelling surgical size is not inherently meant to replicate the tumor’s exact form and proportions, but instead to elucidate the degree of the tissue volume that may be surgically removed in terms of improving patient survival and minimize the risk that subsequent operations will be needed to eliminate all malignant cells entirely. Given the broad range of models of tumor growth, there is no specific rule of thumb about how to select the most suitable model for a particular breast cancer type and whether that would influence its subsequent application in surgery planning. Typically, these models require tumor biology-dependent parametrization, which hardly generalizes to cope with tumor heterogeneity. In addition, the datasets are limited in size, owing to the restricted or expensive measurement methods. We address the shortcomings that incomplete biological specifications, the variety of tumor types, and the limited size of the data bring to existing mechanistic tumor growth models and introduce a Machine Learning model for the PRediction of INdividual breast Cancer Evolution to Surgical Size (PRINCESS). This is a data-driven model based on neural networks capable of unsupervised learning of cancer growth curves. PRINCESS learns the temporal evolution of the tumor along with the underlying distribution of the measurement space. We demonstrate the superior accuracy of PRINCESS, against four typically used tumor growth models, in learning tumor growth curves from a set of four clinical breast cancer datasets. Our experiments show that, without any modification, PRINCESS can accurately predict tumor sizes while being versatile between breast cancer types.

## I. Background

Theoretically, if we would be able to extract cancer’s trajectory from neoplastic cells to metastasis, we could build a model that would anticipate its growth curve, and implicitly tumor size. Of course, we cannot do that due to our incomplete knowledge of tumor biology and the fact that repeated measures of growing tumors are very difficult to obtain. Be it in vitro, in vivo, or through non intrusive imaging, obtaining accurate size measures of a tumor, without influencing the physiology of either tumor or host, is still a challenge. Nevertheless, such an assessment is crucial as it guides cancer therapy. Primarily owing to these shortcomings, researchers investigated ‘‘growth laws” to describe general kinetics of the tumor without regard to some specific form of tumor. These growth laws have a twofold objective: 1) testing growth hypotheses against experimental data and 2) estimating the prior or future course of tumor progression for therapy.

Various patterns of tumor growth were identified experimentally and clinically, and modelled over the years. Among these many ordinary differential equations (ODE) tumor growth models [1] have been proposed and are used to make predictions in cancer treatments planning. In our work, we chose four of the most representative and typically used scalar growth models, namely Logistic, von Bertalanffy, Gompertz and Holling, described in Table I. while such rival canonical tumor growth models are widely used, how to specify which of the models to use on certain types of tumors remains to be determined [2]. Despite their ubiquitous use, in either original form or embedded in more complex models [8], the aforementioned scalar tumor growth models are confined due to: a) the requirement of a precise biological description (i.e. values for *α, β*, λ and *k* correspond to biophysical processes); b) incapacity to describe the diversity of tumor types (i.e. each is priming on a type of tumor), and c) the small amount and irregular sampling of the data (e.g. 15 measurements with at days 6, 10, 13, 17, 19, 21, 24, 28, 31, 34, 38, 41, 45, 48 for breast cancer growth in [9]). As we observe, the need for a Machine Learning system such as PRINCESS is motivated by clinical needs and the peculiarities of tumor growth data. Usually, tumor growth data is small, only a few data points with, typically, days level granularity [10] and irregular spacing among measurements [9]. Moreover, the data has high variability due to: within tumor types specifics, therapy effect on tumor growth [11], and heterogeneous measurement types (e.g. bio-markers, fMRI, fluorescence imaging [12], flow cytometry, or calipers [13]). PRINCESS is a system that attempts to compensate for these shortcomings and exploit a data-driven approach in order to provide precise tumor size estimates.

**TABLE I.** Overview of tumor growth models in our study. Parameters: *N* - cell population size (or volume / mass thorough conversion [7]), *α* - growth rate, *β* - cell death rate, λ - nutrient limited proliferation rate, *k* - carrying capacity of cells.

| Model | Equation |
| --- | --- |
| Logistic [3] | $\frac{dN}{dt} = \alpha N - \beta N^2$ |
| Bertalanffy [4] | $\frac{dN}{dt} = \alpha N^\lambda - \beta N$ |
| Gompertz [5] | $\frac{dN}{dt} = N(\beta - \alpha \ln N)$ |
| Holling [6] | $\frac{dN}{dt} = \frac{\alpha N}{k+N} - \beta N$ |

## II. Materials and methods

In the current section we describe the underlying Machine Learning functionality of PRINCESS along with the datasets used in our experiments.

### A. Introducing PRINCESS

PRINCESS is an unsupervised machine learning system based on Self-Organizing Maps (SOM) [14] and Hebbian Learning (HL) [15] as main ingredients for extracting underlying relations among correlated timeseries describing tumor growth. In order to introduce PRINCESS, we provide a simple example in Figure 1. Here, we consider data from a cubic growth law (3^rd^ powerlaw) describing the impact of sequential cytostatics dose density over a 150 weeks horizon in adjuvant chemotherapy of breast cancer [16]. The two input timeseries (i.e. the number of cancer cells and the irregular measurement index over the weeks) follow a cubic dependency, depicted in Figure 1a.

**Fig. 1.**
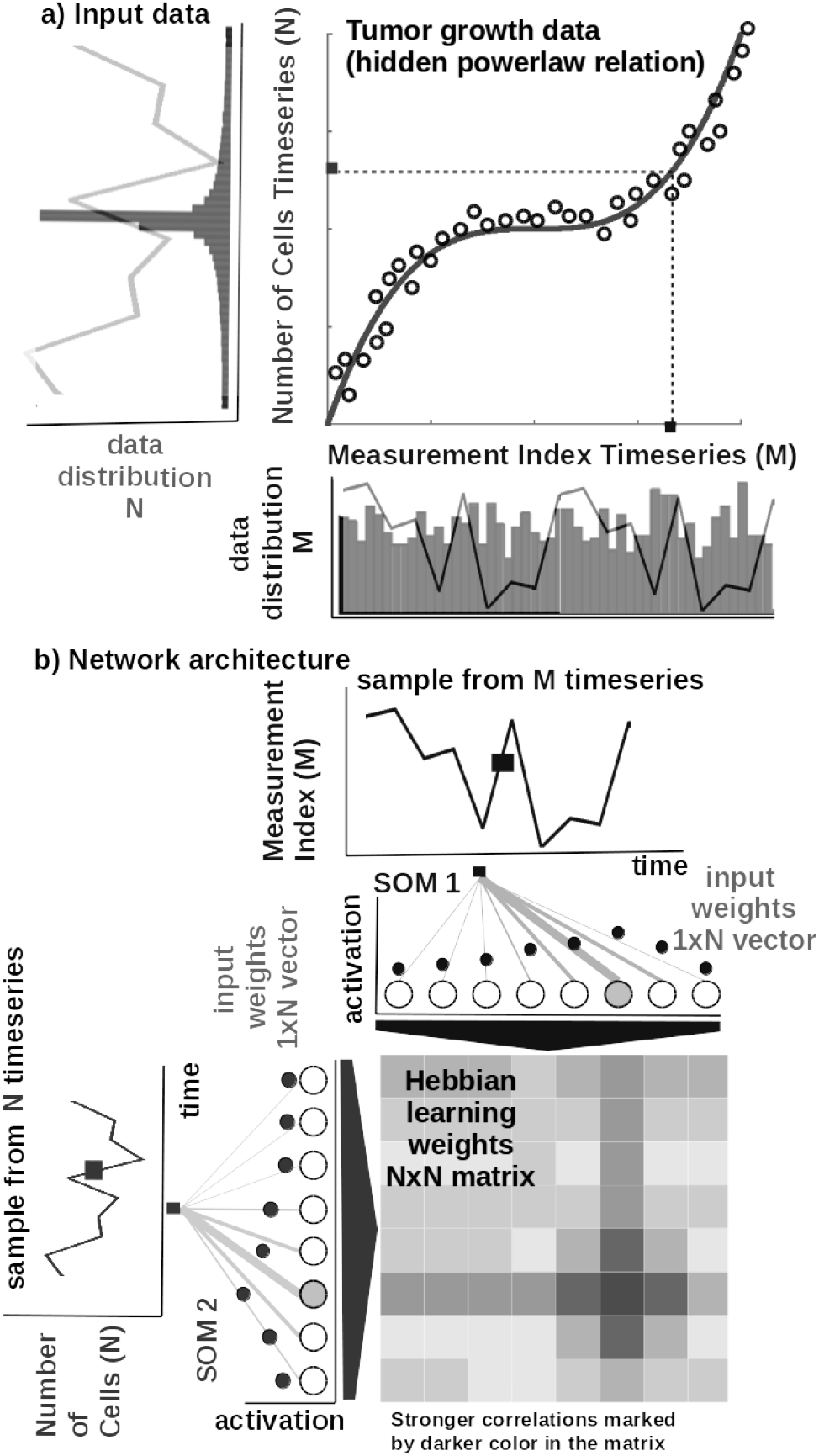
Basic functionality of PRINCESS: a) Tumor growth data resembling a non-linear relation and its distribution - relation is hidden in the timeseries (i.e. number of cells vs. measurement index). Data from [16]. b) Basic architecture of PRINCESS: 1D SOM networks with *N* neurons encoding the timeseries (i.e. number of cells vs. measurement index), and a *NxN* Hebbian connection matrix coupling the two 1D SOMs that will eventually encode the relation between the timeseries, i.e. growth curve.

#### Core model

The input SOMs (i.e. 1D lattice networks with *N* neurons) are responsible to extract the distribution of the incoming timeseries data, depicted in Figure 1a, and encode timeseries samples in a distributed activity pattern, as shown in Figure 1b. This activity pattern is generated such that the closest preferred value of a neuron to the input sample will be strongly activated and will decay, proportional with distance, for neighbouring units. The SOM specialises to represent a certain (preferred) value in the timeseries and learns its sensitivity, by updating its tuning curves shape.

Given an input sample *s^p^*(*k*) from one timeseries at time step *k*, the network computes for each *i*-th neuron in the *p*-th input SOM (with preferred value 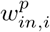 and tuning curve size 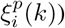) the elicited neural activation as

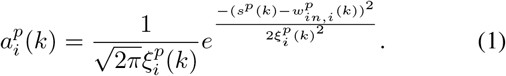

The winning neuron of the *p*-th population, *b^p^*(*k*), is the one which elicits the highest activation given the timeseries sample at time *k*

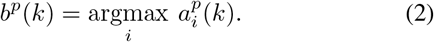

The competition for highest activation in the SOM is followed by cooperation in representing the input space. Hence, given the winning neuron, *b^p^*(*k*), the cooperation kernel,

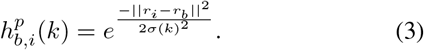

allows neighbouring neurons (i.e. found at position *r_i_* in the network) to precisely represent the input sample given their location in the neighbourhood *σ*(*k*) of the winning neuron. The neighbourhood width *σ*(*k*) decays in time, to avoid twisting effects in the SOM. The cooperation kernel in Equation 3, ensures that specific neurons in the network specialise on different areas in the input space, such that the input weights (i.e. preferred values) of the neurons are pulled closer to the input sample,

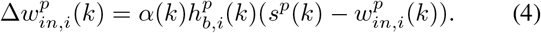

This corresponds to updating the tuning curves width 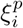 as modulated by the spatial location of the neuron in the network, the distance to the input sample, the cooperation kernel size, and a decaying learning rate *α*(*k*),

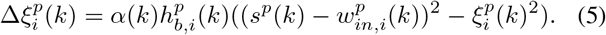

As an illustration of the process, let’s consider learned tuning curves shapes for 5 neurons in the input SOMs (i.e. neurons 1, 6, 13, 40, 45) encoding the breast cancer cubic tumor growth law, depicted in Figure 2. We observe that higher input probability distributions are represented by dense and sharp tuning curves (e.g. neuron 1, 6, 13 in SOM1), whereas lower or uniform probability distributions are represented by more sparse and wide tuning curves (e.g. neuron 40, 45 in SOM1). Using this mechanism, the network optimally allocates resources (i.e. neurons). A higher amount of neurons to areas in the input space, which need a finer resolution; and a lower amount for more coarsely represented areas. Neurons in the two SOMs are then linked by a fully (all-to-all) connected matrix of synaptic connections, where the weights in the matrix are computed using Hebbian learning. The connections between uncorrelated (or weakly correlated) neurons in each population (i.e. *w_cross_*) are suppressed (i.e. darker color) while correlated neurons connections are enhanced (i.e. brighter color), as depicted in Figure 2. Formally, the connection weight 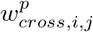 between neurons *i, j* in the different input SOMs are updated with a Hebbian learning rule as follows:

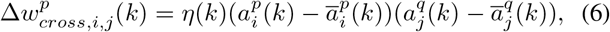

where

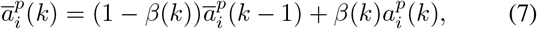

is a “momentum” like exponential decay and *η*(*k*), *β*(*k*) are monotonic (inverse time) decaying functions. Hebbian learning ensures that when neurons fire synchronously their connection strengths increase, whereas if their firing patterns are anticorrelated the weights decrease. The weight matrix encodes the co-activation patterns between the input layers (i.e. SOMs), as shown in Figure 1b, and, eventually, the learned growth law (i.e. relation) given the timeseries, as shown in Figure 2.

**Fig. 2.**
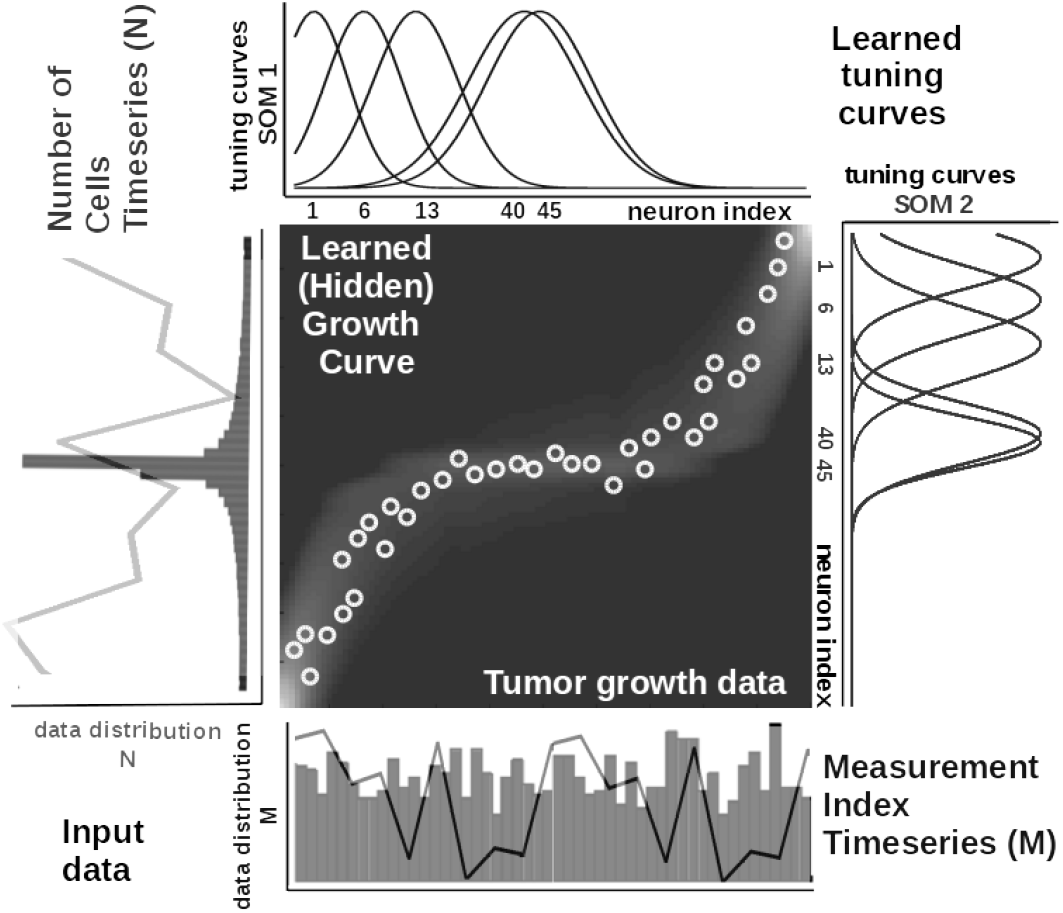
Extracted timeseries relation describing the growth law and data statistics for the data in Figure 1a depicting a cubic breast cancer tumor growth law among number of cells and irregular measurement over 150 weeks, data from [16]. Timeseries overlay on the data distribution and corresponding model encoding tuning curves shapes.

Self-organisation and Hebbian correlation learning processes evolve simultaneously, such that both the representation and the extracted relation are continuously refined, as new samples are presented. This can be observed in the encoding and decoding functions where the input activations are projected though *w_in_* (Equation 1) to the Hebbian matrix and then decoded through *w_cross_*.

#### Parametrization and read-out

In all of our experiments data from tumor growth timeseries is fed to the PRINCESS neural network which encodes each timeseries in the SOMs and learns the underlying relation in the Hebbian matrix. The SOMs neural netwirks are responsible of bringing the timeseries in the same latent representation space where they can interact (i.e. through their internal correlation). In our experiments, each of the SOM has *N* = 100 neurons, the Hebbian connection matrix has size *NxN* and parametrization is done as: *alpha* = [0.01,0.1] decaying, *η* = 0.9, 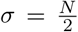 decaying following an inverse time law. We use as decoding mechanism an optimisation method that recovers the real-world value given the self-calculated bounds of the input timeseries. The bounds are obtained as minimum and maximum of a cost function of the distance between the current preferred value of the winning neuron and the input sample at the SOM level.

### B. Datasets

In our experiments we used publicly available tumor growth datasets (see Table II), with real clinical tumor volume measurements, for different cell lines of breast cancer. This choice is to probe and demonstrate transfer capabilities of the models to tumor growth patterns induced by different types of cancer.

**TABLE II.**
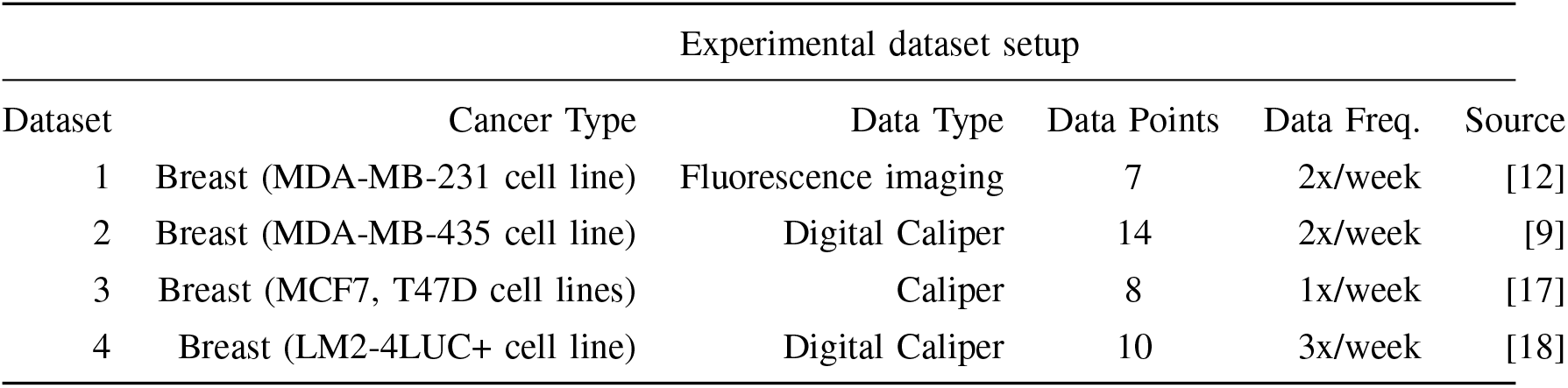
Description of the datasets used in the experiments.

### C. Procedures

In order to reproduce the experiments, the MATLAB^®^ code and copies of all the datasets are available on GITLAB ^1^. Each of the four mechanistic tumor growth models (i.e. Logistic, Bertalanffy, Gompertz, Holling) and PRINCESS were presented the tumor growth data in each of the datasets. When a dataset contained multiple trials, a random one was chosen.

#### Mechanistic models setup

Each of the four tumor growth models was implemented as ordinary differential equation (ODE) and integrated over the dataset length. To provide initial values and the best parameters (i.e. *α, β, λ, k*) for each of the four ODE models a search algorithm was used. Finally, fitting was performed by minimizing the sum of squared residuals (SSR).

#### PRINCESS setup

For PRINCESS the data was normalized before training and de-normalized for the evaluation. The system was comprised of two input SOMs, each with *N* = 50 neurons, encoding the volume data and the irregular sampling time sequence, respectively. Both input density learning and cross-modal learning cycles were bound to 100 epochs.

## III. Results

In the current section we introduce the results, discuss the findings and demonstrate that PRINCESS can provide a precise estimate of surgical size, consistent with measured data in a clinical setup.

### A. Tumor growth curve prediction

In our first batch of experiments, we explore the tumor growth curve prediction of PRINCESS across multiple breast cancer datasets. In all experiments, the four mechanistic models and PRINCESS were evaluated using Sum of Squared Errors (SSE), Root Mean Squared Error (RMSE), and Symmetric Mean Absolute Percentage Error (sMAPE), with definitions in Table III. As our experiments demonstrate (Table IV), PRINCESS obtains superior accuracy in predicting the growth curve of four types of tumors in four cell lines of breast cancer. Such a difference is supported by the fact that PRINCESS is a data-driven Machine Learing model that captures the peculiarities of the data and exploits its statistics to make predictions. The mechanistic models, on the other side, exploit biological knowledge and describe the dynamics of the physical processes but fail in terms of versatility among tumor types. Moreover, due to its learning capabilities PRINCESS excels in capturing the variability and non-uniform data distribution characterising tumor growth data, as shown in the summary statistics overview in Figure 3.

**Fig. 3.**
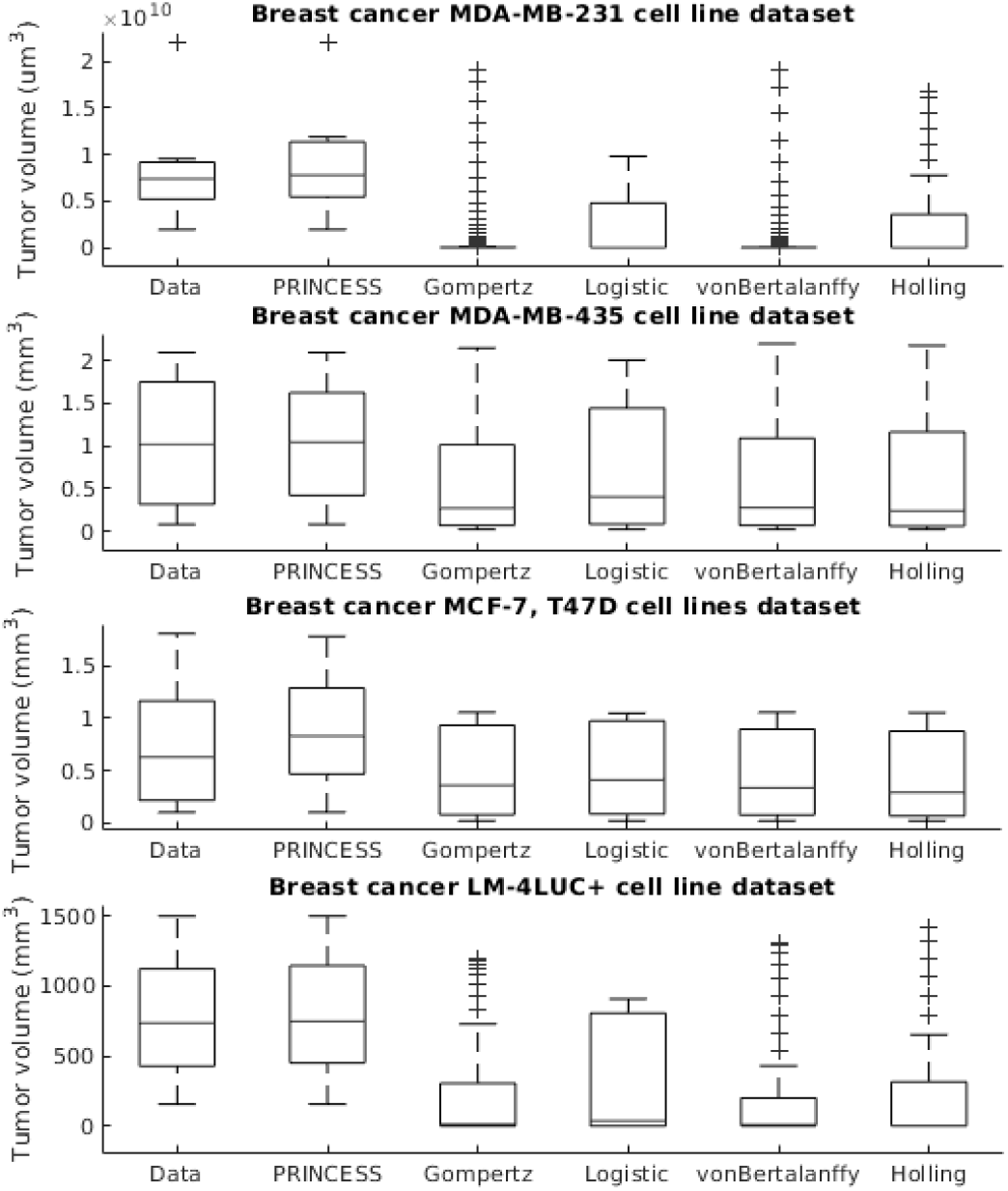
Evaluation of the tumor growth models: summary statistics.

**TABLE III.**
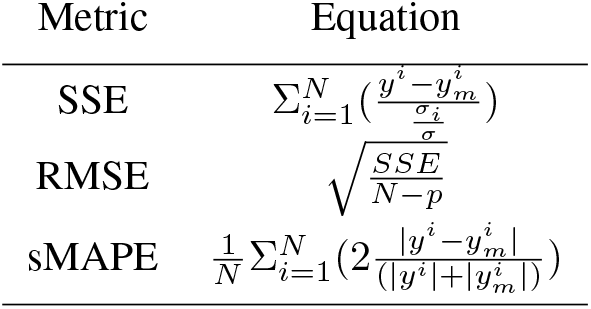
Evaluation metrics for tumor growth models. We consider: *N* - number of measurements, *σ* - standard deviation of data, *p* - number of parameters of the model.

**TABLE IV.** Evaluation of the tumor growth models.

| Evaluation Metrics |  |  |  |
| --- | --- | --- | --- |
| Dataset/Model | SSE | RMSE | sMAPE |
| *Breast<sup>a</sup> cancer [12]* |  |  |  |
| Logistic | 7009.6 | 37.4423 | 1.7088 |
| Bertalanffy | 8004.9 | 44.7350 | 1.7088 |
| Gompertz | 7971.8 | 39.9294 | 1.7088 |
| Holling | 6639.1 | 40.7403 | 1.4855 |
| **PRINCESS** | 119.3 | 4.1285 | 0.0767 |
| *Breast<sup>b</sup> cancer [9]* |  |  |  |
| Logistic | 0.2936 | 0.1713 | 0.1437 |
| Bertalanffy | 0.2315 | 0.1604 | 0.1437 |
| Gompertz | 0.3175 | 0.1782 | 0.1437 |
| Holling | 0.2699 | 0.1732 | 0.1512 |
| **PRINCESS** | 0.0977 | 0.0902 | 0.0763 |
| *Breast<sup>c</sup> cancer [17]* |  |  |  |
| Logistic | 3.0007 | 0.7071 | 1.0606 |
| Bertalanffy | 3.2942 | 0.8116 | 1.0606 |
| Gompertz | 3.1908 | 0.7292 | 1.0606 |
| Holling | 3.4772 | 0.8339 | 1.1288 |
| **PRINCESS** | 0.7668 | 0.3096 | 0.2615 |
| *Breast<sup>d</sup> cancer [18]* |  |  |  |
| Logistic | 45.6031 | 2.3875 | 1.4816 |
| Bertalanffy | 56.0738 | 2.8302 | 1.4816 |
| Gompertz | 53.2428 | 2.5797 | 1.4816 |
| Holling | 54.7189 | 2.7958 | 1.5086 |
| **PRINCESS** | 0.2008 | 0.1417 | 0.0364 |

<sup>a</sup>MDA-MB-231 cell line
<sup>b</sup>MDA-MB-435 cell line
<sup>c</sup>MCF7, T47D cell lines
<sup>d</sup>LM-4LUC+ cell line

### B. Prediction of tumor surgical size

In this section, based on the tumor growth prediction capabilities of PRINCESS, we demonstrate that it can provably predict the surgical size of a tumor from pathology data on an individual patient basis. Based on the tumor growth curves, modelling surgical volume aims to elucidate the extent of the volume of tissue that must be surgically removed in order to (1) increase patient survival and (2) decrease the likelihood that a second or third surgery, or even (3) determine the sequencing with chemotherapy (i.e. adjuvant vs. neo-adjuvant chemotherapy). We assessed this capability by demonstrating that PRINCESS can learn the dependency between histopathological and morphological data, such as nutrient diffusion penetration length within the breast tissue (*L*), ratio of cell apoptosis to proliferation rates (*A*) and radius of the breast tumor (*R*), in an unsupervised manner, from Ductal Carcinoma In Situ (DCIS) data of [19]. More precisely, the authors postulated that the value of R depends upon *A* and *L* following a ‘‘master equation” 8.

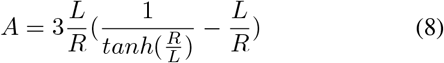

Its predictions, the study shown, are consistent with published findings that nearly 80% of in-situ tumors identified by mammographic screenings. Compared to ground truth, (Equation 8) PRINCESS was capable of extracting an accurate depiction of the growth pattern with *SSE* = 54.216, *RMSE* = 0.4656 and *sMAPE* = 0.5734, respectively. Our belief is that measuring such parameters at the time of initial biopsy, pathologists could use PRINCESS to precisely estimate the tumor size and thus advise the surgeon how much tissue (surgical volume) is to be removed.

## IV. Conclusion

Notwithstanding, tumor growth models have had a profound influence on modern chemotherapy, especially dose management, multi-drug strategies and adjuvant chemotherapy. Yet, the selection of the best model and its parametrization is not always straightforward. To tackle this challenge PRINCESS comes as a support tool that could assist oncologists in extracting tumor growth curves from the data. Using unsupervised learning, PRINCESS overcomes the limitations that incomplete biological descriptions, the diversity of tumor types, and the small size of the data bring on the basic, scalar growth models (e.g. Logistic, Bertalanffy, Gompertz, Holling). PRINCESS exploits the temporal evolution of timeseries of growth data along with its distribution and obtains superior accuracy over basic models in extracting growth curves from different clinical tumor datasets. Without changes either to structure or parameters PRINCESS’s versatility has been demonstrated among different types of breast cancer (i.e. breast MDA-MB-231, MDA-MB-435, LM-4LUC+, MCF7 and T47D cell lines) for predicting tumor growth curves and estimating DCIS surgical size.

**Fig. 4.**
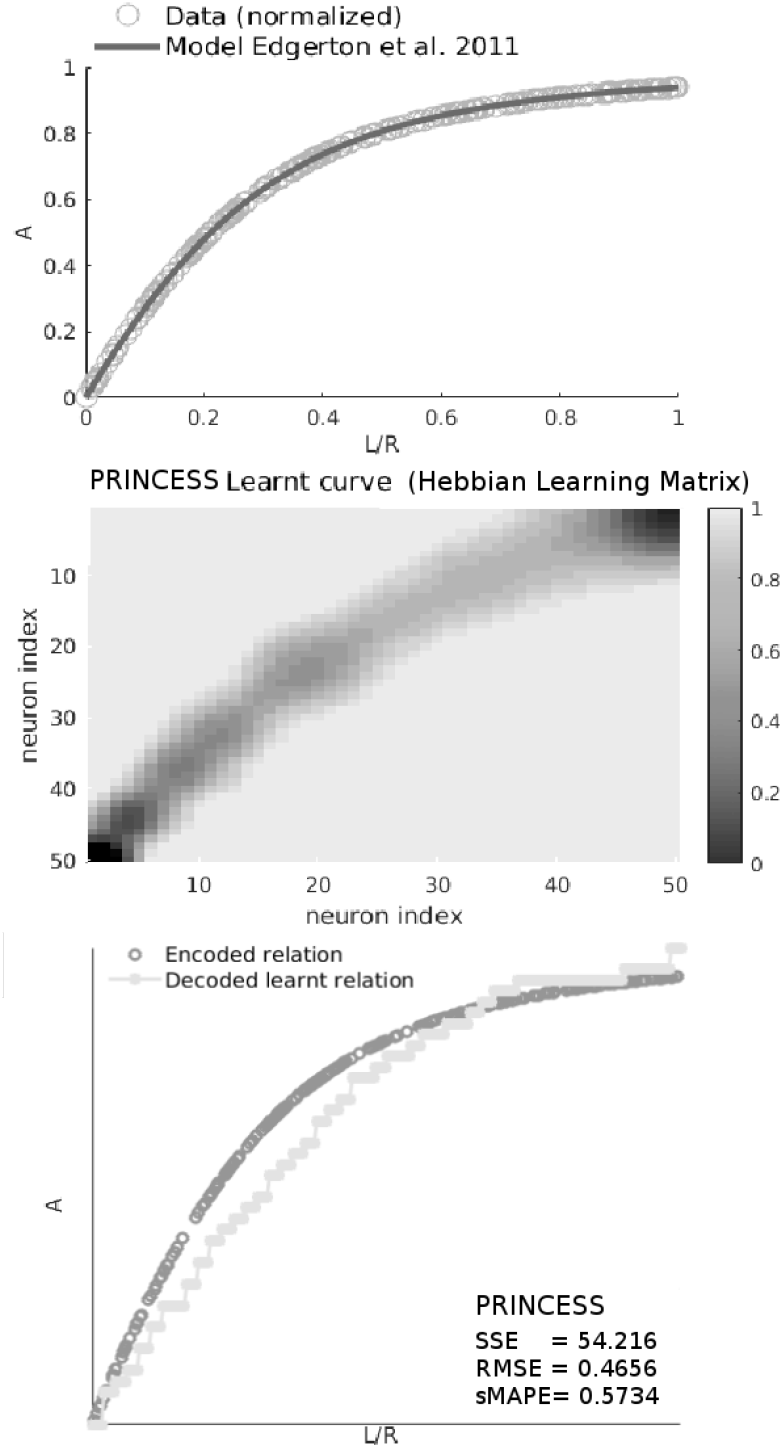
Upper panel: Modelled relation among *A* - the ratio of cell apoptosis to proliferation rates, *L* - the nutrient diffusion penetration length across tumor surgical volume, and *R* - the geometric mean tumor surgical radius, corresponding to 8 on data from [19]. Middle panel: The learnt curve as visible in PRINCESS neural weight matrix (i.e. Hebbian connection matrix). Lower panel: Evaluation of the learnt relation against the analytical (ground truth) model in 8

## Footnotes

1 https://gitlab.com/akii-microlab/cbms2020

